# Screening of additives for droplet qRT-PCR thermocycling enables single influenza A virus genome quantification

**DOI:** 10.1101/2020.04.28.065342

**Authors:** Emma Kate Loveday, Geoffrey K. Zath, Dimitri A. Bikos, Zackary J. Jay, Connie B. Chang

## Abstract

The miniaturization of real time quantitative polymerase chain reaction (qPCR) using drop-based microfluidics, or droplet qPCR, allows for quantification of single nucleic acids. The nucleic acids are compartmentalized into aqueous microdroplets, picoliters in volume, separated by an immiscible oil, and stabilized by a surfactant. In droplet qPCR, accurate data can only be obtained if the drops remain stable to coalescence upon thermocycling and drop contents do not diffuse to neighboring drops. In this work, we present a droplet qRT-PCR assay for quantifying influenza A virus (IAV) following systematic testing of different PCR additives, resulting in the optimal combination of Tween-20 / BSA / betaine to maintain drop stability and limit dye diffusion. We use a standard qPCR machine to generate real time amplification curves of hundreds of thousands of drops and correlate this data with constructed amplification curves obtained from hundreds of drops sampled at various cycle numbers and imaged using epifluorescence microscopy. To demonstrate the utility of our method, we tested a range of *in vitro* transcribed M gene and IAV viral supernatant from infected cells. We directly amplified IAV genomes from infected supernatant without an RNA extraction step. Our droplet qPCR assay enables detection of IAV down to 0.274 cpd, or a single viral genome per drop, establishing the high sensitivity and precision of our method.

## Introduction

The reduction of polymerase chain reaction (PCR) volume using drop-based microfluidics has emerged in the past decade as a powerful method for increasing the speed and sensitivity of nucleic acid detection.^1-7^ Such droplet PCR applications involve the amplification of nucleic acids that are compartmentalized into picoliter-sized aqueous microdroplets, separated by an immiscible oil phase, and allow for high-throughput analysis with reduced reagent and sample consumption.^3-5, 8, 9^ Two methods of performing droplet PCR include droplet digital PCR (ddPCR)^1, 2^ and quantitative PCR (qPCR).^3-5, 10, 11^ The ddPCR method is most frequently used to quantify rare nucleic acid species via limiting dilution, in which drops containing nucleic acids and ddPCR mix are diluted to such a limit that each drop contains either zero or at least one nucleic acid copy.^1, 7, 12^ In this case, endpoint fluorescent drop measurements provide a digital readout where positive drops containing template are bright and negative drops without template are dim. The fraction of positive drops is used to calculate starting nucleic acid concentration. Currently, the majority of ddPCR applications rely on the use of proprietary reagent mixtures for droplet production and a commercial device to measure and quantify drop fluorescence.^1, 2^ Droplet qPCR assays utilize a similar approach in which a sample and PCR mix are combined together within picoliter-sized drops and amplification during thermocycling is measured with a fluorescent probe in real time as opposed to an endpoint measurement.^3-5, 10, 11^ Droplet qPCR provides a method for analysis and quantification of nucleic acid and allows for characterization of large heterogenous populations in a high throughput manner.^1, 5, 6, 8^

Drops used for PCR applications can be created using biocompatible fluorinated oils with perfluorinated surfactants stabilizing the water-in-oil drops. A commonly used perfluorinated surfactant is the non-ionic, tri-block copolymer perfluoropolyether-polyethylene glycol-perfluoropolyether (PEG-PFPE_2_).^13^ The PEG group extends into the aqueous drop, making the surfactant biocompatible when present among cells, nucleic acids, and proteins.^13^ However, drops made with PEG-PFPE_2_ have been shown to destabilize under increased thermal conditions above 60 °C, a temperature significantly exceeded during PCR thermocycling.^14, 15^ In addition to stability issues, increased temperature during thermocycling promotes transport between drops and results in small molecules, such as the qPCR probe and reference dye, to diffuse between the drops.^16-18^ As qPCR assays rely on fluorescence signals from probes and reference dyes, drop stability and dye retention during thermocycling is critical for the detection and precise quantification of genomic content in drops.^19^ To maintain drop stability and dye retention, droplet PCR assays stabilized with PEG-PFPE_2_ use high surfactant concentrations (2-5 wt %)^8, 15, 20^, small drop sizes (<40 μm diameter), and/or incorporated additives. The most common additives used for droplet PCR assays are polyethylene glycol (PEG), Tween-20, bovine serum albumin (BSA), and betaine; yet systematic testing of the effect of these additives is critically lacking and, to our knowledge, has not been previously reported. Systematic testing would provide an optimized set of additives for droplet qPCR by improving drop stability during thermocycling and limiting transport of small molecules between drops.

Here, we investigate a range of additives for optimizing droplet quantitative reverse transcriptase PCR (qRT-PCR) and enabling quantification of influenza A virus (IAV) in 100 μm diameter drops. Additive combinations of PEG, Tween-20, BSA, and betaine were screened for their ability to stabilize drops from coalescence and limit diffusion between neighboring drops. Together with 3 wt% (w/w) of PEG-PFPE_2_ surfactant^13^ in the oil phase, an additive combination of 1% w/v Tween-20, 0.8 µg/µl BSA, and 1 M betaine in a standard qRT-PCR mixture was found to be optimal for stabilizing drops and limiting dye diffusion under qPCR thermocycling between 60 °C and 95 °C. Using Tween-20 / BSA / betaine as additives, five orders of magnitude of *in vitro* transcribed viral RNA was resolved between drops ranging from 10^−1^ to 10^4^ copies per drop (cpd). Drops containing low (10^1^ cpd) and high (10^4^ cpd) viral RNA concentrations were thermocycled using a standard qPCR machine and the qRT-PCR amplification curves were comparable to constructed curves obtained from drops sampled at various cycles using epifluorescence microscopy. The optimized droplet qRT-PCR assay was used to quantify the number of A/California/07/2009 (H1N1) IAV genomes present in the supernatant from infected A549 cells. At 24 hours post infection (hpi), we prepared six dilutions of supernatant from infected cells to determine our detection limit for live virus. Using ddPCR analysis, we were able to quantify IAV RNA directly from infected cells across four orders of magnitude. We find that our limit of detection is at a dilution that corresponds to 0.274 cpd, demonstrating that we are able to detect down to a single viral genome in a drop. In addition, we amplified viral RNA from supernatant collected directed from infected cells without an RNA extraction step. This direct PCR method will have impact in reducing the number of steps for future work performing droplet qPCR of viruses.

## Materials and Methods

### TaqMan qRT-PCR Reactions

The sequences of qRT-PCR amplification primers for the IAV Matrix gene (M gene) as described by Shu et al. were as follows: Matrix gene forward primer 5’-GACCRATCCTGTCACCTCTGAC-3’, Matrix gene reverse primer 5’-AGGGCATTCTGGACAAATCGTCTA-3’.^21^ The sequence of Matrix gene TaqMan probe was: 5’-/FAM/TGCAGTCCTCGCTCACTGGGCACG/BHQ1/-3’. Primers and probe were ordered from Eurofins Operon and were prepared as 100 µM stocks. The working stocks of the primers were 25 µM with a final concentration of 400 nM. The working stock of the probe was 10 µM with a final concentration of 200 nM. Samples were amplified using a SuperScript III Platinum One-Step qRT-PCR kit (Invitrogen 11732-020) with a final reaction volume of 25 µL. Each reaction mix contained 0.05 µM ROX reference dye, 2 mM MgSO_4_ and 0.32 U/µL SUPERase RNase Inhibitor (Invitrogen AM2694), and 2.5 µL of RNA or supernatant. Tested additives were added to the qRT-PCR reaction mix at the following concentrations: 1% w/v Tween-20 (Calbiochem 655204-100mL), 0.8 µg/µL BSA (Fisher BP675-1), 2.5% w/v PEG-6K (Acros Organics 192280010), and 1 M betaine (Sigma B0300-1VL). Thermocycling was performed in a qRT-PCR machine (Quantstudio 3, Applied Biosystems) with the following cycling conditions: 1 cycle for 30 min at 60 °C, 1 cycle for 2 min at 95 °C, and 40 cycles between 15 s at 95 °C and 1 min at 60 °C.

### *In vitro* Transcribed RNA

To quantify the amount of viral RNA in each drop, standard curves were generated for each run using serial dilutions of *in vitro* transcribed IAV segment 7 (Matrix gene).^22, 23^ The nucleic acid copy number in each reaction or drop was extrapolated from the standard curve. To generate the *in vitro* transcribed RNA, a gBlock containing a T7 promoter (underlined), forward and reverse primer sites (italicized), and probe sequence (bold) for the Matrix gene was ordered from IDT: 5’-GTC<u>TAATACGACTCACTATAG</u>*GACCAATCCTGTCACCTCTGAC* **TGCAGTCCTCGCTCACTGGGCACG** TGCTTCATCGCGAACTGCTTCGCGGATGCCATCGTCATGGCCACGAGGATATGTAAG AGT *TAGACGATTTGTCCAGAATGCCCT*-3’. The Matrix gene was transcribed *in vitro* using a MEGAscript™ T7 RNA Synthesis Kit (Ambion, AM1333) and purified over a GE Illustra Sephadex G-50 NICK column. The RNA was then DNase treated and precipitated with ammonium acetate/ethanol according to the manufacturer’s instructions. RNA concentration was quantified using a NanoDrop spectrophotometer and used to determine the copy number per µL.

### Microfluidic device fabrication

Microfluidic devices for making 50 μm and 100 μm diameter drops were fabricated by patterning SU-8 photoresist (Microchem SU-8 3050) on silicon wafers (University Wafer, ID# 447, test grade) using standard photolithography techniques.^24^ Polydimethylsiloxane (PDMS) (Sylgard 184) at 10:1 mass ratio of polymer to cross-linking agent is poured onto the patterned device master molds. Air was purged from the uncured PDMS by placing the filled mold in a vacuum chamber for at least 1 h. The PDMS was cured in an oven at 55 °C for 24 h and then removed from the mold with a scalpel. Ports were punched into the PDMS slab with a 0.75 mm diameter biopsy punch (EMS Rapid-Core, Electron Microscopy Sciences). The PDMS slab was bonded to a 2-in by 3-in glass slide (VWR micro slides, cat. #48382-179) after plasma treatment (Harrick Plasma PDC-001) for 60 s at high power (30 W) and 700 mTorr oxygen pressure. The drop making devices were made hydrophobic by flowing Aquapel (Pittsburgh Glass Works) through the channels, followed by blowing the channels with air filtered through a GVS ABLUO™ 25 mm 0.2-μm filter (Fisher Scientific) before baking the devices in an oven at 55 °C for 1 h.

### Drop encapsulation

Flow-focusing drop making devices were used to make 50 µm diameter drops for the drop stability and PCR dilution experiments and 100 µm diameter drops for infected supernatant experiments.^25^ The dispersed phase consisted of qRT-PCR mix containing a range of concentrations (10^3^ to 10^8^ copies/µL) of *in vitro* transcribed M gene RNA and different combinations of additives (no additives, Tween-20 only, BSA only, PEG-6K only, betaine only, Tween-20/BSA, Tween-20/PEG-6K and Tween20/BSA/betaine with concentrations as previously described). The continuous phase consisted of a 3 wt% (w/w) solution of PEG-PFPE_2_-based surfactant (RAN Biotechnologies, 008-FluoroSurfactant) in fluorinated HFE7500 oil (3 M). The dispersed and continuous phases were loaded in 1-mL syringes and injected into the drop making microfluidic devices at a flow rate of 800 µL/h and 1600 μL/h, respectively, using syringe pumps (New Era NE-1000) controlled by a custom LabVIEW (2015) program to generate drops. A total volume of 60 µL composed of approximately 20 µL of drops and 40 µL of oil were collected in 100 µL PCR tubes (Applied Biosystems). After the drops were collected, the PCR tubes were promptly placed in the qRT-PCR machine (Applied Biosystems QS3) and thermocycled.

### Endpoint brightfield and fluorescence imaging of drops containing PCR additives combinations

Thermocycled drops were imaged by injecting 20 μL of drops into a microfluidic device with a wide inlet channel large enough to capture a full field of view (FOV) of drops with a 10x objective. The channel height of the device was 50 μm. Brightfield and fluorescence images (Texas RED (TXRED, ex. 540-580 nm, em. 592-669 nm)) of the drops were captured on an inverted epifluorescence microscope (Leica DMi8) with a 10x objective (Leica, NA 0.32). The Leica Application Suite X was used for image acquisition. For each of the eight additive conditions, drops were imaged from three PCR tubes and three FOV per tube (>100 drops per FOV) were captured.

### Analysis of endpoint brightfield and fluorescence imaging of drops containing combinations of PCR additives

Drops imaged on the epifluorescence microscope were processed with a custom MATLAB (R2019a) script to measure drop diameter and ROX fluorescence intensity. Measurements of drop diameters *D* from at least 270 drops per condition were made following thermocycling. All drop measurements were made on the TXRED channel of the acquired images. ROX fluorescence intensities *I* were normalized by subtracting the background fluorescence *I*_B_. Background fluorescence was set as the average pixel value measured from a small (100 px^2^) background region in the middle of the image using FIJI.^26^ The device used for imaging had a channel height of 50 µm, thereby compressing drops larger than 50 μm in diameter and skewing larger drop diameter measurements. Thus, drop diameters greater than this height were adjusted by estimating the compressed drops as oblate spheroids and then using the oblate spheroid volume to calculate the equivalent sphere diameter using the equation 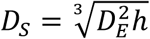 where *D*_S_ is the diameter of the equivalent sphere, *D*_E_ is the measured diameter of a compressed drop, and *h* is the channel height of 50 μm. Normalization of measured *D* from an initial drop diameter *D*_0_ was determined with the equation (*D* − *D*_0_)/*D*_0_. The initial *D*_0_ is the average drop diameter measured from drops without PCR additives upon exit from a drop making device as shown in **Fig. S1**. *D*_0_ was used to normalize the diameter of drops that contained different additives following thermocycling.

### Brightfield and fluorescence imaging of drops during thermocycling

For drops imaged before, during, and after thermocycling at each specified cycle, the qPCR machine was paused and one PCR tube was removed to sample drops. Drops were imaged using epifluorescence microscopy to quantify fluorescence from the FAM (6-carboxyfluorescein) dye-labeled TaqMan probe and ROX (6-carboxy-X-rhodamine) reference dye. Five FOV were captured of drops from each tube at each cycle on the epifluorescence microscope for an average of 650 drops after pooling from all FOV (≈0.02% of the drops contained in the PCR tube). The fluorescence intensities of individual drops were measured using a custom MATLAB script (R2019a).

### Analysis of brightfield and fluorescence imaging of drops during thermocycling

Drops imaged on the epifluorescence microscope were processed with a custom MATLAB (R2019a) script to measure FAM fluorescence intensity. The FAM fluorescence intensities measured during qRT-PCR were normalized as Δ*R*_*N*_/*R*_*N*,0_, where the qPCR normalized reporter value *R*_N_ is defined as *R*_*N*_ = *I*_*FAM*_/*I*_*RON*_, Δ*R*_*N*_ = *R*_*N*_ − *R*_*N,baseline*_, *I*_FAM_ is the FAM intensity at each cycle, *I*_ROX_ is the ROX intensity at each cycle, *R*_N,baseline_ is the average *R*_N_ value of the first three cycles, and *R*_N,0_ is the initial *R*_N_ value.

### Virus strains and cell lines

The IAV virus strain used in the present study was A/California/07/2009 (H1N1) (kindly provided by Dr. Christopher Brooke). Viral A/California/07/2009 (H1N1) stocks were propagated and titered on Madin-Darby Canine Kidney (MDCK) cells (kindly provided by Dr. Agnieszka Rynda-Apple). MDCK cells were propagated in DMEM (Corning) media supplemented with 10% fetal bovine serum (HyClone) and 1x Penicillin/Streptomycin (Fisher Scientific). All experimental infections were performed on human alveolar epithelial A549 cells (kindly provided by Dr. Christopher Brooke). A549 cells were propagated in Hams F-12 media (Corning) supplemented with 10% fetal bovine serum and 1x Penicillin/Streptomycin.

### Virus infections

A549 cells were seeded into 6-well plates with 1 x 10^6^ cells per well. The cells were infected with the H1N1 virus at a multiplicity of infection (MOI) of 0.1 in infection media consisting of Hams F-12 supplemented with 1-mM HEPES (HyClone), 1x Penicillin/Streptomycin (Fisher Scientific) and 0.1% BSA (MP Biomedical). Infection with the H1N1 virus was performed in the presence of 1 µg of TPCK (tolylsulfonyl phenylalanyl chloromethyl ketone)-trypsin/ml (Worthington Biomedical).^27^ Briefly, cells were washed with 1x Phosphate Buffered Saline (PBS) (Corning) and then incubated with virus that was diluted in infection media for 1 h. The inoculum was then removed and replaced with fresh infection media and supernatant was collected at 24 hpi.

## Results and Discussion

### Screening of qRT-PCR additives to stabilize drops and reduce dye diffusion during thermocycling

To probe how different PCR additives impact drop stability and dye diffusion during thermocycling, we measured the effect of Tween-20, BSA, PEG-6K, and betaine individually and in different combinations in a droplet qRT-PCR assay with 3 wt% (w/w) of PEG-PFPE_2_ surfactant. To analyze drop stability, drop diameters *D* were measured post thermocycling from epifluorescence images. To quantify dye diffusion, fluorescence intensities of a thermally stable reference dye (ROX) that is present in all droplet qPCR assays was measured post thermocycling from epifluorescence images. All of the droplet qRT-PCR assays were optimized for TaqMan-based chemistry to detect and quantify *in vitro* transcribed M gene RNA from IAV. The M gene is a standard target for IAV detection due to its conserved nature across different IAV species.^21, 28^ Drops were produced from a solution of qRT-PCR reaction master mix, probe and primers, enzymes and 10^7^ copies/µL (∼170 cpd) of M gene template. This represents the qRT-PCR reaction mixture with no additives. To test different PCR additives, the qRT-PCR reaction mixture was supplemented with each of the following: BSA, PEG-6K, betaine, Tween-20, Tween-20 / PEG-6K, Tween-20 / BSA, or Tween-20 / BSA / betaine. PEG enhances PCR reactions due to its ability to act as a macromolecular crowding agent, thereby increasing DNA polymerase activity.^29^ In drops, PEG has been shown to improve the stability of drops containing high salt content which can occur in standard PCR reaction mix.^30-32^ BSA is a common PCR additive and has been shown previously to limit dye diffusion between drops.^33, 34^ Tween-20 has been utilized as an additional surfactant^8, 35, 36^ and acts to reduce surface tension in drops while betaine improves PCR amplification of GC rich regions by reducing the formation of nucleic acid secondary structure.^37-39^

The drops containing 10^7^ copies/µL of M gene template were thermocycled on a standard qPCR machine with a 30-minute reverse transcriptase step followed by 40 cycles of PCR. Drops were imaged on an epifluorescence microscope following thermocycling in order to assess drop stability from coalescence and retention of the ROX dye. The images of drops that contained no additives, BSA alone, PEG-6K alone, or betaine alone show extensive coalescence with multiple different drop sizes in each image (**Fig. 1A**). Diameters *D* were compared to the initial drop diameter *D*_0_ before thermocycling of drops with no PCR additives. Normalized diameters (*D*-*D*_0_)/*D*_0_ closer to zero indicate less change in drop diameter after thermocycling. The (*D*-*D*_0_)/*D*_0_ of the various additive combinations in **Fig. 1B** demonstrate that drops containing no additives, or only a single additive of BSA, PEG-6K, or betaine, had high deviation from zero, a wide distribution of diameters, and a large number of outliers as represented by individual dots. The addition of Tween-20 alone to the PCR reaction mix resulted in a tight distribution of drop diameters. However, (*D*-*D*_0_)/*D*_0_ was below zero with a value of -0.12, indicating that the Tween-20 drops decreased in size from their initial diameters (**Fig. 1B, red box**). As the addition of Tween-20 decreased drop coalescence upon thermocycling, we proceeded to test different combinations of the additives with Tween-20. Tween-20 was combined with PEG-6K, BSA and BSA / betaine. The individual additives PEG-6K, BSA, and betaine were chosen as they were utilized in prior droplet PCR assays.^20, 35, 40, 41^ By adding Tween-20 to each of these additives, (*D*-*D*_0_)/*D*_0_ decreased below zero and were found to have lower coefficients of variation (CV) than with no additives and with each of the additives alone without Tween-20 (**Fig. 1B**). The measured diameters indicate that the addition of Tween-20 greatly decreased both drop coalescence and CV, although epifluorescence images show that Tween-20 / PEG-6K and Tween-20 / BSA combinations still contained a number of large drops, indicating that the problem of coalescence was not fully resolved. The combination of Tween-20 / BSA / betaine drops resulted in (*D*-*D*_0_)/*D*_0_ closest to zero at -0.04 with very few outliers, indicating these drops were the most stable following thermocycling (**Fig. 1B, orange box**).

**Figure 1:**
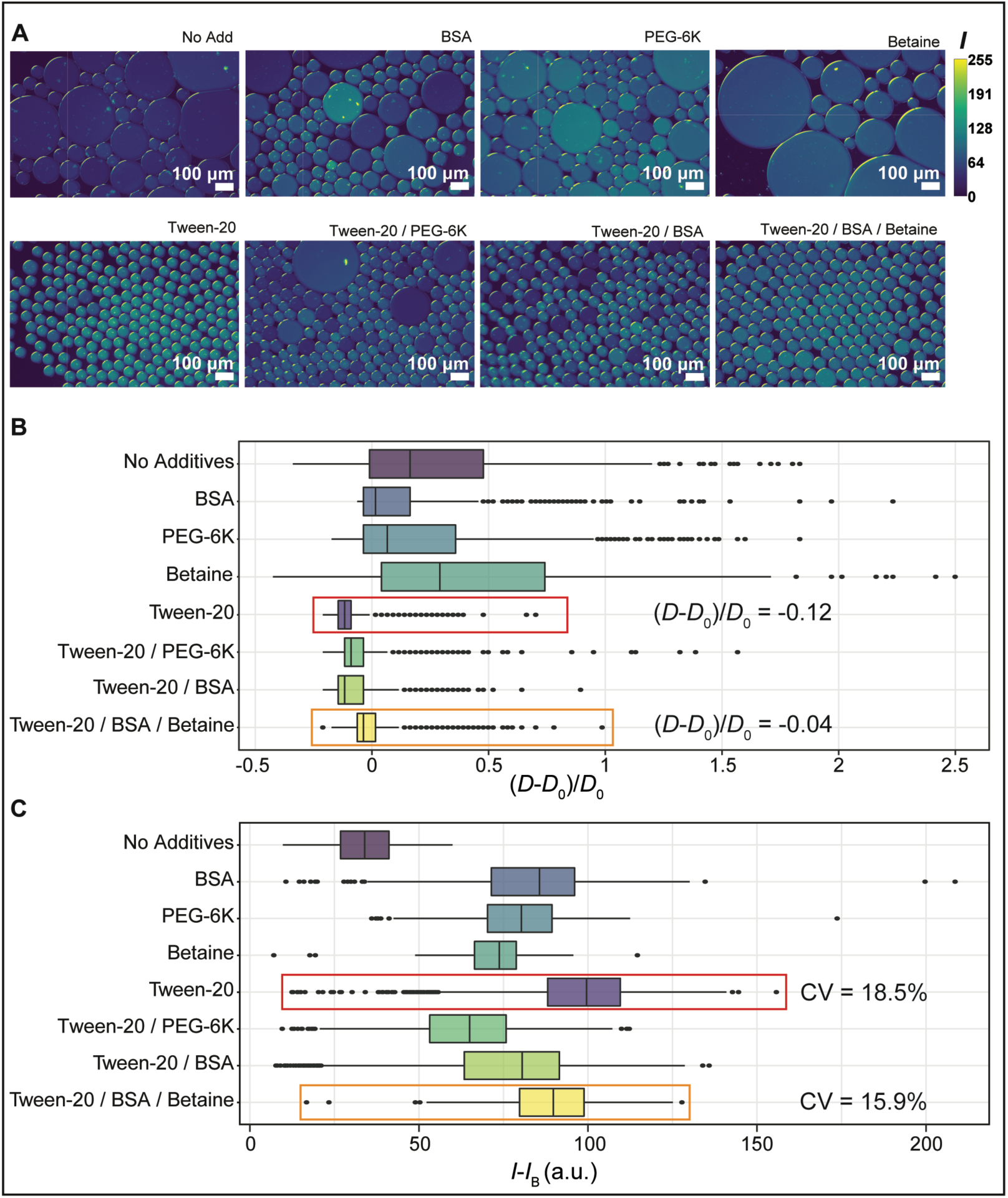
Addition of additives during PCR thermal cycling. Tween-20, BSA, PEG-6K, and betaine were investigated as additives individually and in various combinations to improve drop stability and prevent dye diffusion. **(A)** Drop stability and dye diffusion were measured from epifluorescence images of thermocycled drops with representative images shown of drops with no additives, BSA, PEG-6K, Betaine, Tween-20, Tween-20 / PEG-6K, Tween-20 / BSA, or Tween-20 / BSA / betaine. Fluorescence intensity *I* is quantified from pixel values ranging from 0 to 255. Boxplots of **(B)** normalized diameter (*D*-*D*_0_)/*D*_0_ and **(C)** average ROX fluorescence intensity of drops *I-I*_*B*_ following qPCR with different additives. Drops are distributed by quartile, with the vertical line within the box representing the median value and the boxes represent the middle quartile (25%-75% distribution). The lines represent drops in the upper and the lower 25% of the distribution. Dots represent individual drops and extreme values that are either 1.5X larger or smaller than the interquartile range.

In addition to drop stability, we also investigated the retention of ROX reference dye within drops after thermocycling. The fluorescence intensity *I* of ROX in each of the epifluorescence images was normalized by subtracting the background signal *I*_*B*._ The normalized ROX fluorescence intensity of each drop, *I-I*_*B*_, was measured to quantify diffusion of the dye from the drops (**Fig. 1C**). A lower value of *I-I*_*B*_ indicates that the ROX dye diffused out of the drops and not between neighboring drops. To verify this, we measured the background fluorescence of the oil surrounding the drops and found that the background fluorescence intensity was highest for the no additive condition (**Fig. S2**), indicating that the additives tested here had a positive effect on ROX dye retention in drops during thermocycling.

A narrow ROX *I-I*_*B*_ within drops (smaller CV) is ideal for normalizing the TaqMan probe signal that is produced during thermocycling. In addition, a high *I-I*_*B*_ indicates that the dye is not diffusing out of the drops. Thus, we tested drop additives to look for a narrow distribution CV and overall high ROX fluorescence intensity (**Fig. 1C**). With no additives, *I-I*_*B*_ of the ROX dye was 33.6 a.u., the lowest of all eight conditions tested, and indicates a loss of ROX dye from the drops during thermocycling. Based on our data, Tween-20 and the Tween-20 / BSA / betaine combination maintained the best drop stability post thermocycling. In terms of ROX dye retention, Tween-20 had the highest *I-I*_*B*_ of 99.6 a.u. (**Fig. 1C, red box**) compared to Tween-20 / BSA / betaine with a *I-I*_*B*_ of 89.7 a.u. (**Fig. 1C, orange box**). However, the Tween-20 / BSA / betaine condition resulted in a smaller CV (15.9%) of *I-I*_*B*_ than the Tween-20 condition (18.5%). Based on our results, droplet qRT-PCR reactions that contain an additive combination of Tween-20 / BSA / betaine have the lowest change in drop diameters following thermocycling with relatively high overall ROX fluorescence intensity and the lowest CV of the fluorescence intensity. Numbers of drops measured and statistical information from **Fig. 1B-C** can be found in **Table S1**.

### Dilution series demonstrates high efficiency for droplet qRT-PCR

To evaluate the reaction efficiency of our droplet qRT-PCR reaction with the optimized additives Tween-20 / BSA / betaine, we performed a dilution series of RNA in drops and as bulk reactions. PCR is based on the principle that amplification begins in an exponential growth phase at early thermocycles before the reaction products become exhausted. This exponential phase corresponds to a reaction efficiency of 100%, where all DNA is multiplied by a factor of two at the completion of every cycle. Desired amplification efficiencies should fall between 90-110%.^42, 43^ PCR efficiencies that fall outside of this range will limit dynamic range and sensitivity. A dilution series of a known quantity of target RNA or DNA is required to create a standard curve to relate cycle threshold (C_t_) values to the common logarithm (log_10_) of the RNA or DNA concentration. When a standard curve has a slope of -3.33, it indicates that the PCR reaction efficiency is 100%. To perform a standard curve in drops, *in vitro* transcribed IAV M gene RNA was added as template to the optimized qRT-PCR reaction mix with additives Tween-20 / BSA / betaine. Serial RNA dilutions of a range of 10^4^ to 10^9^ copies/μL in bulk were generated, corresponding to a range of 10^−1^ to 10^4^ cpd. The amplification curves obtained using qPCR in bulk **(Fig. S3)** follow the same trend as in drops **(Fig. 2A)**. The C_t_ values for the dilution series in drops and in bulk were plotted to generate standard curves for both reactions **(Fig. 2B)**. The slopes of the curves yield a droplet reaction efficiency of 90.3% compared to a bulk reaction efficiency of 98.9%, with both falling within the desired 90-110% range (**Fig. 2B**). We hypothesize that the lower efficiency in the droplet reactions compared to bulk is due to sequestration of reagents within the drops, resulting in a slightly reduced ability to fully amplify the DNA over 40 cycles before exhaustion of required reaction reagents.

**Figure 2:**
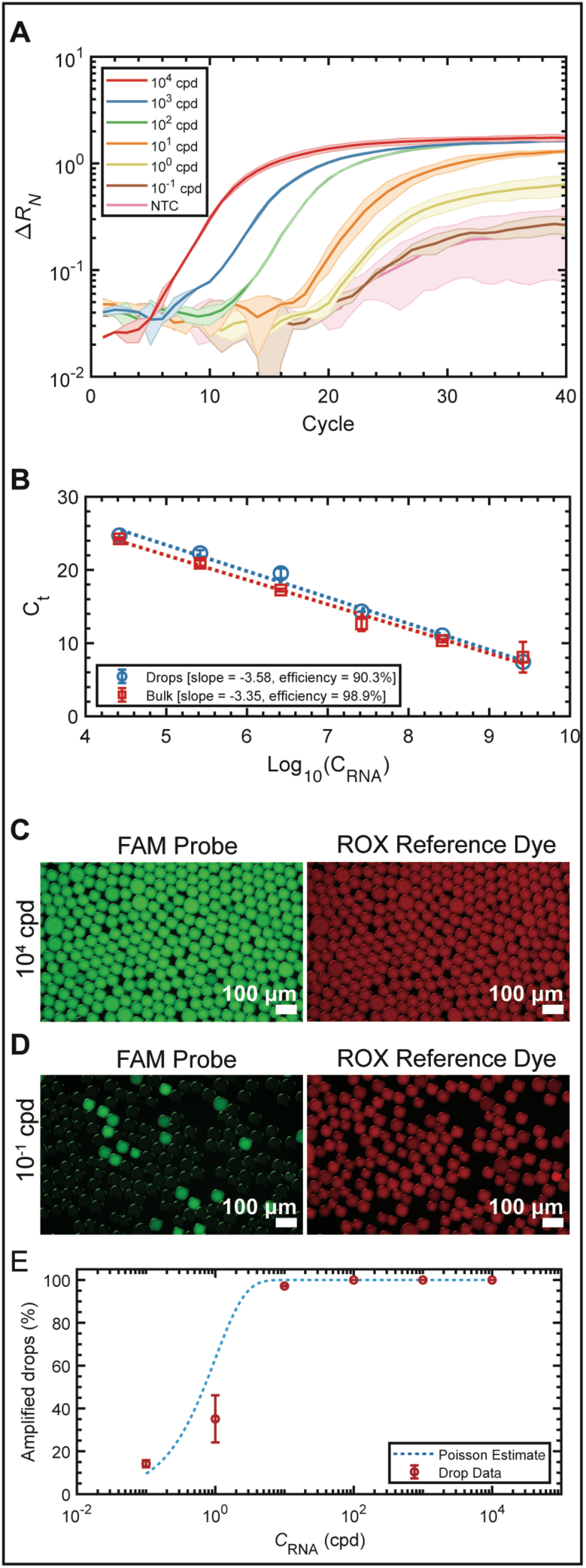
qRT-PCR dilution series of *in vitro* transcribed IAV M gene in drops. **(A)** Amplification curves of six 10-fold dilutions of M gene RNA amplified in drops ranging from 10^−1^ to 10^4^ copies of RNA per drop (cpd). **(B)** C_t_ standard curves for the bulk and drop amplification curves. The calculated qRT-PCR reaction efficiency was 90.3% for the drop dilution series and 98.9% for the bulk dilutions series, which falls within the desired range (90% to 110%). Representative epifluorescence images of the FAM and corresponding ROX channels of drops containing **(C)** 10^4^ cpd and **(D)** 10^−1^ cpd after 40 thermocycles. **(E)** Droplet digital PCR (ddPCR) analysis displaying the percentage of amplified drops as a function of RNA cpd. The total percentage of amplifying bright drops (red circles) increases as a function of RNA concentration and closely follows the Poisson estimate (blue dotted line).

An end-point measurement of the drop fluorescence can be performed either with epifluorescence microscopy or custom flow-based detection methods.^1, 2, 24^ Here, we used epifluorescence microscopy. Drops from the dilution series in **Fig 2A** were imaged on an epifluorescence microscope after 40 PCR cycles. Representative images of FAM and ROX signals in drops containing 10^4^ **(Fig. 2C)** and 10^−1^ **(Fig. 2D)** RNA cpd are shown. As expected, the 10^4^ cpd case resulted in all bright drops due to the high concentration of RNA amplified in each drop, while the 10^−1^ cpd case resulted in a combination of bright and dark drops due to the limiting dilution of RNA. We performed ddPCR analysis by counting the number of bright versus dark drops from the endpoint reactions of the dilution series. Poisson statistics was used to fit the fraction of positive drops from the dilution series images, following **Eq. 1**:

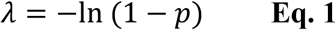

where *λ* is the average copy numbers of target RNA and *p* is the fraction of positive drops.^1^ The percentage of amplified drops demonstrated good agreement with Poisson statistics **(Fig. 2E, Table S2)**. At 10^−1^ cpd and 10^0^ cpd, there were 14.2% and 38.1% amplified drops, respectively, while at higher concentrations (10^1^ through 10^4^ cpd) the number of amplified drops reached 100%, following the Poisson estimate for calculating RNA concentration through ddPCR. Our results demonstrate that our optimized qRT-PCR reaction mix maintained drop stability and prevented dye transport between drops, allowing us to normalize the fluorescent signal from the TaqMan FAM probe in amplified drops. In addition, our droplet qRT-PCR exhibited a high PCR reaction efficiency of 90.3%, which was within the desired range of 90-110% for qPCR reactions.

### Generation of droplet PCR amplification curves from direct qRT-PCR thermocycling compared to epifluorescence microscopy

Concentration of nucleic acid present within a sample can be determined using qRT-PCR. In the prior section, we introduced a novel method for testing qRT-PCR additives to obtain amplification curves across five orders of magnitude of M gene RNA in drops. Here, we performed a high-resolution investigation of two concentrations of M gene in drops undergoing real-time PCR amplification. We studied drops containing low (10^1^ cpd) and high (10^4^ cpd) viral RNA by sampling drops at various cycle numbers on the qRT-PCR machine and imaging under epifluorescence microscopy. We obtained real-time PCR amplification curves from the thermocycled drops in the qPCR machine and compared these curves to constructed amplification curves arising from hundreds of individual drops imaged using epifluorescence microcopy.

Amplifying RNA within drops on a standard qPCR machine generated amplification curves of all the drops contained within the sample tube, approximately 300,000 drops. The qPCR machine tracks fluorescence amplification of all drops in the tube at each cycle number between 0 to 40, which we call continuous data. We sampled hundreds of randomly sampled drops imaged on an epifluorescence microscope over multiple thermocycles, which we call discontinuous data. This allowed us to construct amplification curves using microscopy that resembled the real-time continuous measurements performed on the standard qPCR machine. Thermocycling drops within the qPCR machine without interruption generated a continuous amplification curve **(Fig. 3A, green solid line)** and is representative of a bulk sample **(Fig. S3)**. By contrast, a discontinuous amplification curve was produced by imaging drops from individual tubes at specific cycles on an epifluorescence microscope **(Fig. 3A, blue dashed line)**. Drops from the low RNA loading condition were analyzed across eight cycles (cycle 1, 20, 22, 24, 26, 28, 30, and 40) **(Fig. 3A).** Cycle 1 provides a baseline fluorescence measurement. At cycle 20, the amplification curve begins to increase exponentially. Drops are then imaged after every two cycles from cycle 20 to 30 to capture the exponential region of the amplification curve. A final endpoint measurement is taken at cycle 40. The normalized fluorescence data from the epifluorescence images **(Fig. 3A, blue dashed line)** are plotted for each cycle measured and produce a curve similar to that of the qPCR machine **(Fig. 3A, green solid line)**, validating that an amplification curve can be constructed from the discontinuous epifluorescence data. The distributions of drop fluorescence from epifluorescence images at cycles 1, 20, 22, 24, 26, 28, 30, and 40 for the low RNA loading condition are presented in **Fig. 3B**. As amplification starts to increase at cycle 20, the distribution of drops shifts to the right as the fluorescence increases.

**Figure 3:**
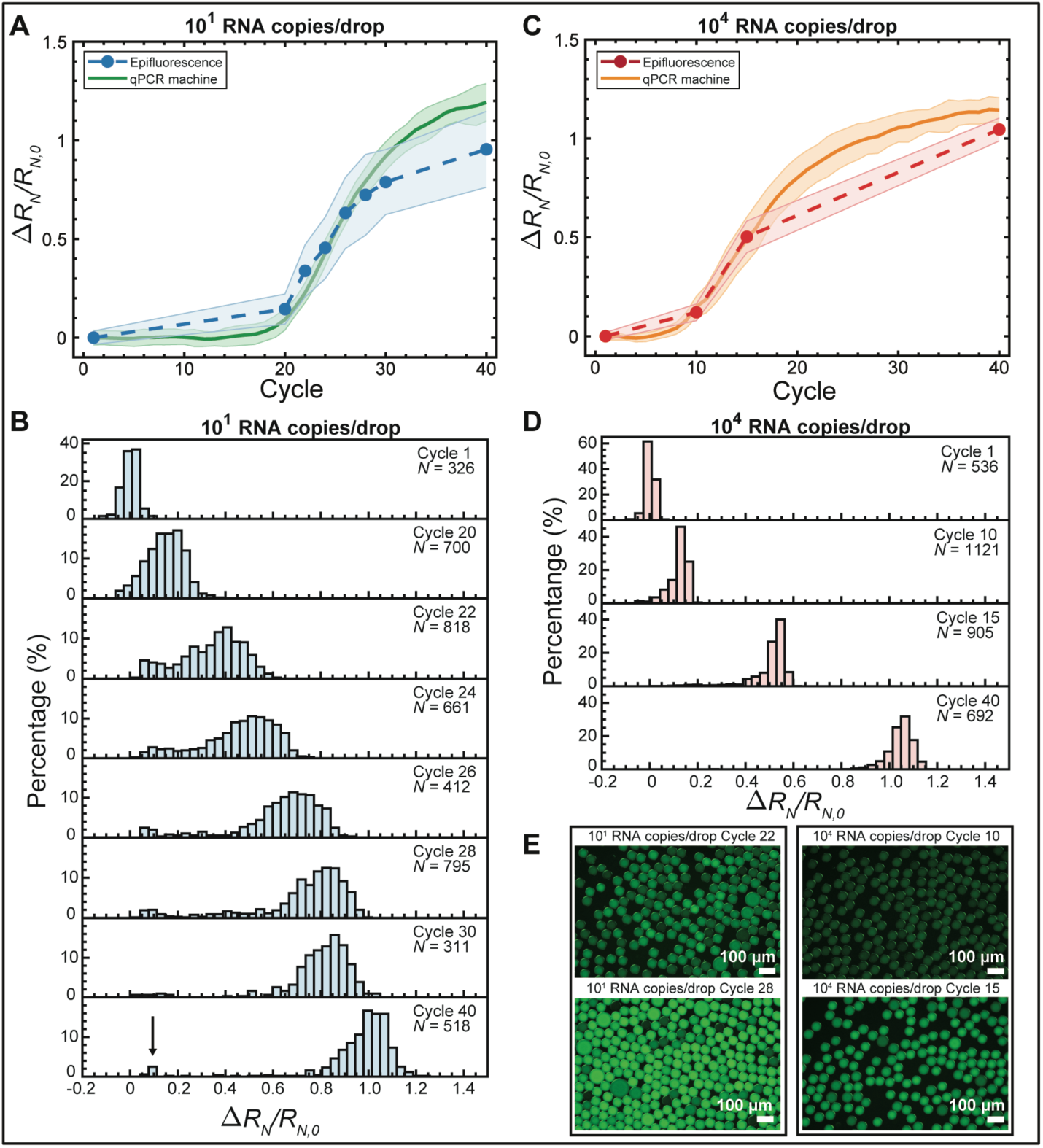
Comparison of droplet qRT-PCR amplification curves generated using a standard qPCR machine and epifluorescence imaging. Drops containing **(A)** 10^1^ M gene RNA cpd were thermocycled using qPCR. The solid green line represents real-time continuous fluorescence measurements of drops at each cycle as determined by qPCR, whereas the blue dashed line represents fluorescence measurements of sampled individual drops at cycle numbers 1, 20, 22, 24, 26, 28, 30, and 40 as determined by epifluorescence microscopy. Shaded error bars represent one standard deviation. **(B)** Histograms of individual drop fluorescence values from the epifluorescence images for 10^1^ M gene RNA cpd show an increase in the fluorescence intensity from cycle 20 to 30. Cycle 1 and 40 represent pre- and post-thermocycling. *N* represents the number of drops analyzed from multiple epifluorescence images for each cycle. The arrow in cycle 40 represents unamplified drops present in our low RNA loading condition. **(C)** 10^4^ M gene RNA cpd were thermocycled using qPCR. The solid orange line represents continuous fluorescence measurements of drops at each cycle as determined by qPCR, whereas the red dashed line represents fluorescence measurements of individual drops at cycle numbers 1, 10, 15, and 40 as determined by epifluorescence microscopy. Shaded error bars represent one standard deviation. **(D)** Histograms of individual drop fluorescence values from the epifluorescence images for 10^4^ M gene RNA cpd show an increase in the fluorescence intensity from cycle 15 to 40. Cycle 1 and 40 represent pre- and post-thermocycling. *N* represents the number of drops analyzed from multiple epifluorescence images for each cycle. **(E)** Changes in fluorescence as cycle numbers increase are observed from representative epifluorescence images of 10^1^ or 10^4^ M gene RNA cpd at cycles 22 and 28 and cycles 10 and 15, respectively.

From cycle 20 to 30, we observed that a majority of the drops began to amplify while a small population did not **(Fig. 3B, arrow, cycle 40)**, forming a bimodal distribution. We hypothesize that this small population of non-amplifying drops did not contain template RNA. This would indicate that the 1.7 x 10^1^ cpd loading was an overestimate as empty drops are not expected at this concentration. In comparison, the high RNA loading condition (1.7 x 10^4^ cpd) should result in every drop containing RNA, with all drops amplifying in unison, and should not result in a bimodal distribution as observed with the low RNA loading condition. To test this, drops containing 1.7 x 10^4^ RNA cpd were imaged at 0, 10, 15 and 40 cycles **(Fig. 3C)**. As with the low RNA loading condition, the normalized fluorescence data from the high RNA loading condition epifluorescence images produced a curve (**Fig. 3C, red dashed line**) similar to that of the qPCR machine (**Fig. 3C, orange line**). The distributions of drop fluorescence at each cycle for the high RNA loading condition are presented in **Fig 3D**.

As discrete particles such as RNA are encapsulated into drops following a Poisson distribution,^24^ we predicted the high RNA loading condition (1.7 x 10^4^ cpd) would create a tight distribution (CV = 0.77%) while the low RNA loading condition (1.7 x 10^1^ cpd) would create a wide distribution (CV = 24.3%) of RNA between drops, respectively. The CV is calculated as 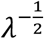, as defined by the Poisson distribution, where *λ* is the RNA loading in cpd. The high RNA loading condition resulted in all drops amplifying as predicted. In addition, the fluorescence intensities of the high loading RNA condition showed tighter distributions at all cycles compared to the amplified drops in the low RNA loading conditions. Representative images of drops for both 1.7 x 10^1^ and 1.7 x 10^4^ cpd at early and late cycles provide a visual reference for the distributions in the histograms **(Fig. 3E)**. The similar trend of drop measurements between the qPCR machine and the epifluorescence images demonstrates that RNA concentrations in individual drops can be extrapolated from real-time PCR amplification curves produced by either standard qPCR thermocycling or epifluorescence microscopy. These methods can therefore be utilized to determine nucleic acid concentrations from a large population of drops without the need for custom flow-based methods.^1, 2, 24^

### qRT-PCR amplification and quantification of live influenza A virus

Following successful amplification of *in vitro* transcribed RNA, we applied our optimized droplet qRT-PCR assay to quantify the number of IAV genomes present in the supernatant from infected cells using ddPCR analysis. We infected A549 cells, a human alveolar epithelial cell line, with the A/California/07/2009 (H1N1) IAV at a low MOI of 0.1. At 24 h post infection, we prepared six dilutions of supernatant from infected cells starting with undiluted, or 10^0^ dilution, down to a 10^−5^ dilution. Supernatant from mock infected cells was included as a control to determine the level of background amplification and the detection limit for live virus. Infected supernatant dilutions and undiluted supernatant from mock infected cells were added to the optimized qRT-PCR master mix and drops were produced, thermocycled, and analyzed on the epifluorescence microscope. In this case, the virus was heat lysed during the reverse transcription step at 60 °C for 30 min and then amplified without an RNA extraction step. This direct PCR method allowed us to perform ddPCR directly from the supernatant of infected cells to quantify the amount of RNA present. We measured the fraction of positive drops to total drops (*N*_+_/*N*_total_), also defined as *p* in **Eq. 1** and plotted this as a function of dilution factor **(Fig. 4A, blue solid line)**. A Poisson fit with an R^2^ value of 0.987 was used to estimate the average number of IAV genomes per drop at each dilution (**Fig. 4A, green dashed line, Table S2**). The undiluted supernatant was estimated to contain 175 cpd, corresponding to 3.34 x 10^6^ copies/μL. We compared this value to the measured concentration of the undiluted supernatant obtained using qRT-PCR. The C_t_ value of the undiluted supernatant resulted in a concentration of 1.96 x 10^6^ copies/μL, based upon the standard curve from **Fig. 2A**, in agreement with our Poisson estimate. The supernatant dilutions **(Fig. 4A, solid blue line)** followed the Poisson fit down to the 10^−3^ dilution (0.175 cpd). The background amplification level of mock infected cells was 0.082 cpd **(Fig. 4A, red dashed line)**. Representative images are shown and demonstrate that undiluted supernatant results in every drop amplifying after 40 cycles **(Fig. 4B)**. In comparison, a 10^−2^ dilution of supernatant, which we estimated to contain 1.75 cpd, results in a majority of drops amplifying (77%) with a few that remain dark **(Fig. 4C)**. The lowest 10^−5^ dilution of supernatant from infected cells is similar to supernatant from mock infected cells and is therefore representative of background amplification **(Fig. 4D and E)**. Our limit of detection was reached at a 10^−3^ dilution level, in which *N*_+_/*N*_total_ was measured to be 0.274 cpd. This demonstrates the ability of our droplet qPCR assay to detect viral RNA down to a single cpd. Thus, we demonstrate the ability to amplify and quantify IAV RNA directly from infected cells across four orders of magnitude, down to single viral genome copies encapsulated within drops.

**Figure 4:**
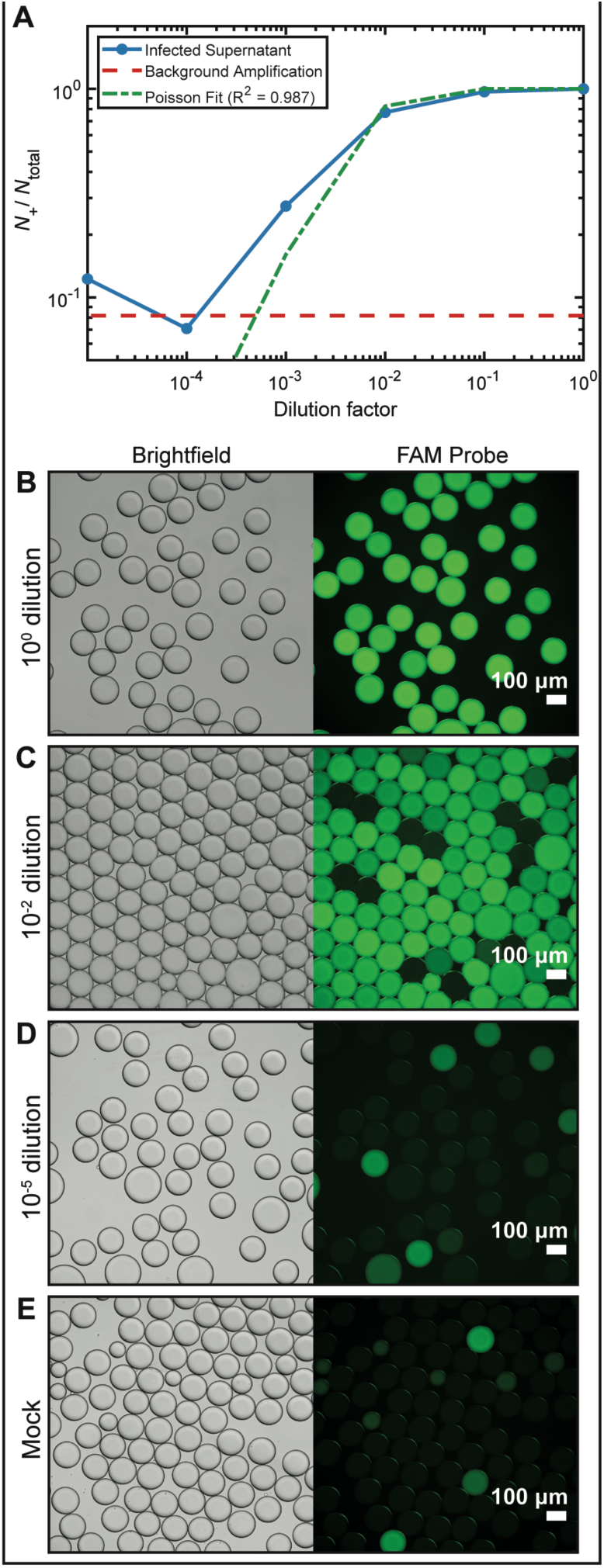
IAV amplified using droplet qRT-PCR method. **(A)** Supernatant from infected cells was diluted and combined in drops with our optimized PCR mixture. The fraction of amplified drops is measured as the fraction of the number of fluorescent drops (*N*_+_) and the total number of drops (*N*_total_). A ddPCR analysis of end point epifluorescence imaging shows the dilution series of viral RNA in drops (solid blue line) coincides with the Poisson fit (dashed green line). Detection of viral RNA was achieved over four orders of magnitude before reaching the level of background amplification as determined using supernatant from mock infected cells (dashed red line). **(B-E)** Representative epifluorescence images of thermocycled drops containing various dilutions of viral supernatant from infected cells, including a mock sample with no virus.

## Conclusions

In this work, we present a droplet qRT-PCR assay for quantifying IAV following systematic testing of different PCR additives, resulting in the optimal combination of Tween-20 / BSA / betaine to maintain drop stability and limit dye diffusion. We demonstrate, for the first time, the ability to use a standard qPCR machine to generate real time amplification curves of hundreds of thousands of drops. We correlate this data with hundreds of drops sampled using epifluorescence microscopy. Measurements of real-time PCR amplification in drops using a standard qPCR machine or epifluorescence microscope can eliminate the need for complicated custom microfluidic thermocycling devices^3-5^. As qPCR machines and epifluorescence microscopes are common instruments in many laboratories, this method can be used to systematically test droplet qPCR additives for quantifying genomes within a large population of drops.

We tested a wide concentration range of both *in vitro* transcribed viral RNA and viral supernatant from infected cells in drops. Our optimized droplet qPCR assay enabled quantification of 10^−1^ to 10^4^ cpd of *in vitro* transcribed IAV M gene. To further demonstrate the utility of our method, we quantified viral IAV genomes from infected cells. Using ddPCR, we measured IAV RNA over four orders of magnitude down to 0.274 cpd, or a single viral genome per drop. This demonstrates the high sensitivity and precision of our droplet qPCR assay.

Importantly, our work establishes the ability to directly amplify viral RNA without the need for RNA extraction and with very little reagent, which is advantageous in challenging resource-limited situations such as the COVID-19 pandemic. We believe our findings will have great impact in future studies for accurately quantifying viral genomes using droplet qPCR.

## Acknowledgements

This work was supported by Defense Advanced Research Projects Agency (DARPA) grant W911NF-17-2-0034.

## Supplemental

**Table S1:** Supplemental data table for Fig. 1 B-C.

| Additive | $(D-D_0)/D_0$<br>mean | $(D-D_0)/D_0$<br>standard<br>deviation | $(D-D_0)/D_0$<br>median | $D$<br>CV | $(I-I_B)$<br>mean | $(I-I_B)$<br>standard<br>deviation | $(I-I_B)$<br>median | $(I-I_B)$<br>CV | $N_{drops}$ |
| --- | --- | --- | --- | --- | --- | --- | --- | --- | --- |
| No Additives | 0.31 | 0.43 | 0.16 | 33% | 33.6 | 10.4 | 34.0 | 30.9% | 405 |
| BSA | 0.12 | 0.28 | 0.01 | 25% | 83.6 | 20.1 | 85.7 | 24.0% | 915 |
| PEG-6K | 0.25 | 0.40 | 0.09 | 32% | 79.2 | 15.0 | 80.0 | 19.0% | 622 |
| Betaine | 0.46 | 0.60 | 0.29 | 41% | 72.6 | 10.9 | 73.8 | 15.1% | 270 |
| Tween-20 | -0.10 | 0.08 | -0.12 | 9% | 97.0 | 17.9 | 99.6 | 18.5% | 1948 |
| Tween-20 / PEG-6K | -0.05 | 0.14 | -0.09 | 15% | 63.9 | 16.9 | 65.3 | 26.4% | 1534 |
| Tween-20 / BSA | -0.07 | 0.11 | -0.12 | 12% | 76.2 | 23.5 | 80.5 | 30.8% | 1460 |
| Tween-20 / BSA / Betaine | -0.01 | 0.12 | -0.04 | 12% | 88.9 | 14.1 | 89.7 | 15.9% | 1155 |

**Table S2:** Supplemental data table for ddPCR analysis Fig. 2E and 4A.

| Dilution Factor | $N_+/N_{total}$<br>(*M* gene standards) | $N_+/N_{total}$<br>(infected supernatant) |
| --- | --- | --- |
| $10^0$ | 1 | 1 |
| $10^{-1}$ | 1 | 0.968 |
| $10^{-2}$ | 1 | 0.771 |
| $10^{-3}$ | 0.972 | 0.274 |
| $10^{-4}$ | 0.351 | 0.07 |
| $10^{-5}$ | 0.142 | 0.123 |
| Negative Control | 0.065 | 0.082 |

**Figure S1:**
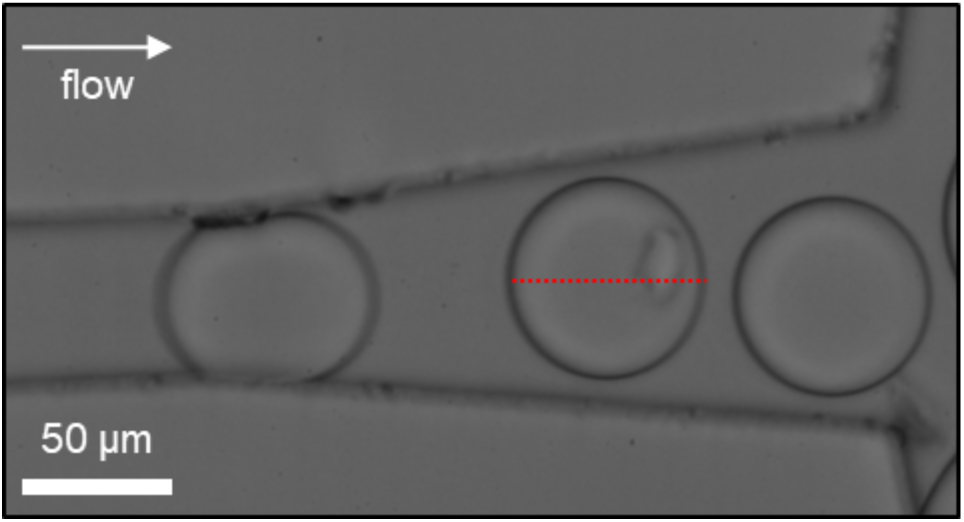
Drops at the exit of a drop making device. Initial drop diameter (*D*_*0*_) is measured as indicated by the dotted red line.

**Figure S2:**
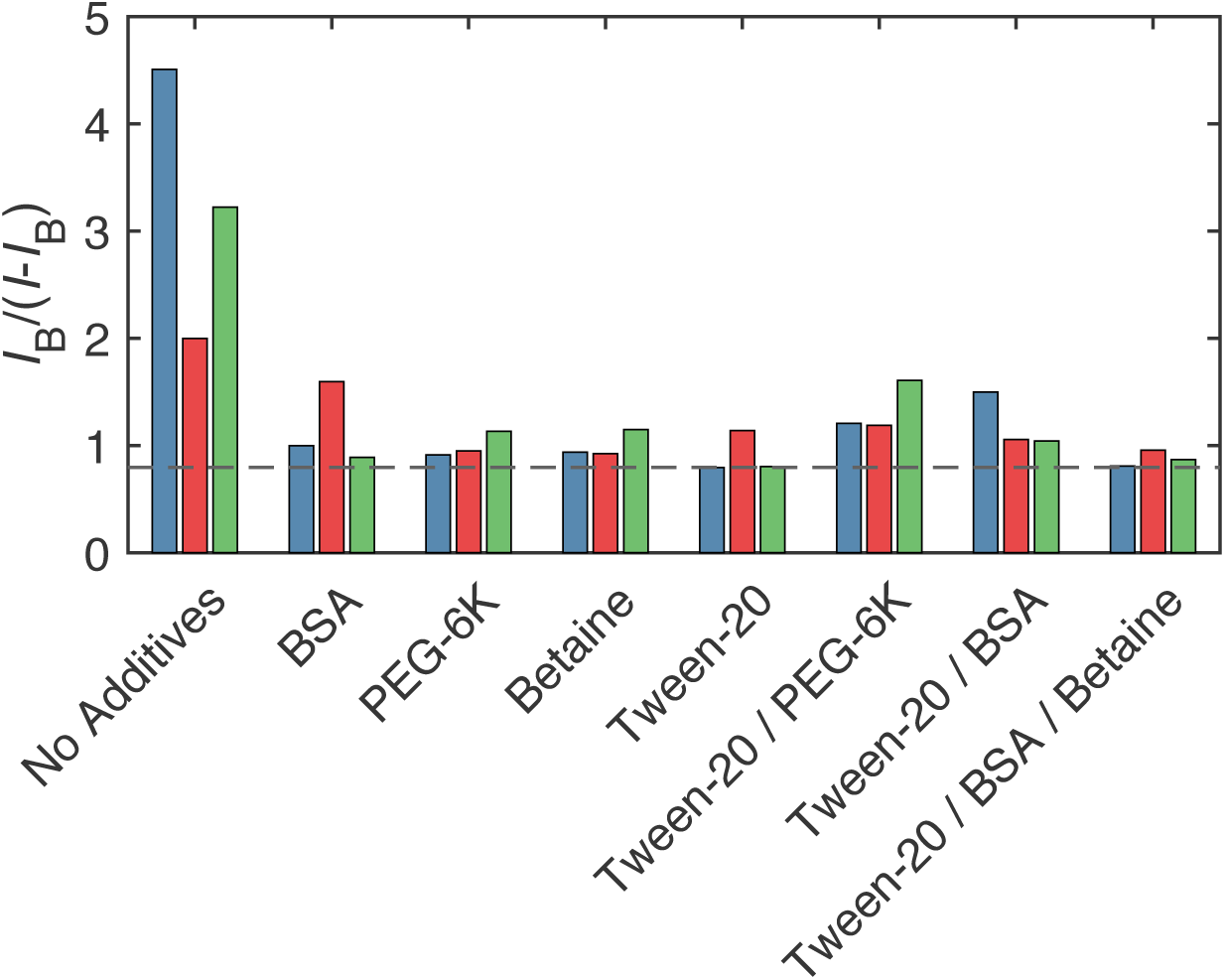
Mean normalized background fluorescence intensity 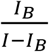 for each PCR additive condition. Colored bars for each additive represent pooled image data from each qPCR tube sampled. The dashed line is used to show that the Tween-20 and Tween-20 / BSA / betaine additive conditions had the lowest normalized background, indicating the best retention of ROX dye in the drops during thermocycling.

**Figure S3:**
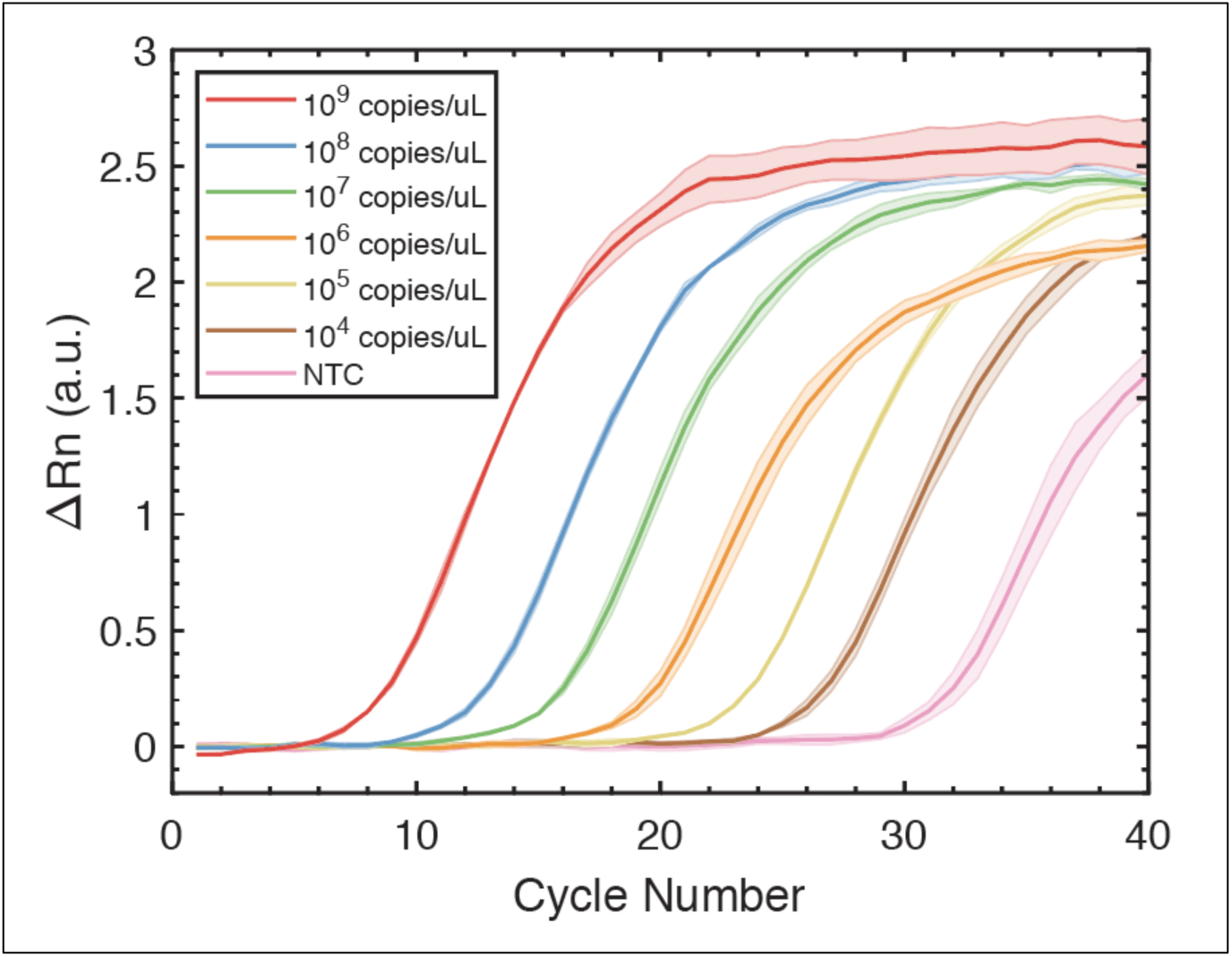
Amplification curves of 6 different dilutions of M gene RNA from bulk samples run on a QuantStudio 3 Real-Time PCR System.

